# vcfpp: a C++ API for rapid processing of the Variant Call Format

**DOI:** 10.1101/2023.10.12.555914

**Authors:** Zilong Li

## Abstract

Given the widespread use of the variant call format (VCF/BCF) coupled with continuous surge in big data, there remains a perpetual demand for fast and flexible methods to manipulate these comprehensive formats across various programming languages. Many bioinformatic tools were developed in C++ to ensure high performance and modern C++ standards offer an ever expanding libraries to ease program development. This work presents vcfpp, a C++ API of HTSlib in a single file, providing an intuitive interface to manipulate VCF/BCF files rapidly and safely, in addition to being portable. Moreover, this work introduces the vcfppR package to demonstrate the development of a high performance R package with vcfpp, allowing for rapid and straightforward variants analyses. In the benchmarking, with the compressed VCF of 3202 samples and one million variants as input, the dynamic script using vcfppR is only 1.3*×* slower than its compiled C++ counterpart vcfpp, whereas the Python API cyvcf2 is 1.9*×* slower when streaming a variant analysis with little memory. Lastly, in a two-step setting where the whole VCF content is loaded first, vcfppR demonstrates a 101*×* speed improvement over vcfR and even more folds than data.table in processing genotypes.

## Introduction

Computational biologists have made numerous efforts and contributions to facilitate analyses of genomic variants. The variant call format (VCF) has become the standard for storing genetic variant information consisting of detailed specifications (Danecek et al. 2011). With big data on the rise, the binary variant call format (BCF) was later designed to query and store large datasets efficiently. The C API of HTSlib (Bonfield et al. 2021) provides a full set of functionalities to manipulate the VCF/BCF for both compressed and uncompressed files. Given that the C API is challenging for less proficient programmers to use, APIs derived from other languages have been created to fill in the gap. Existing popular libraries include vcfR (Knaus et al. 2017) for R, cyvcf2 (Pedersen et al. 2017) for Python, hts-nim (Pedersen et al. 2018) for Nim and vcflib (Garrison et al. 2022) for C++. All are valuable to each respective community, but not without their disadvantages. In particular, vcflib is both an API and a large collection of command line tools, with the primary pitfall being that it does not support the BCF format. It is noteworthy that many methods written in C++ designed for large sample size can not input the compressed VCF or BCF, as in the case of Syllbable-PBWT (Wang et al. 2023). The motivation behind vcfpp is to offer full functionalities as HTSlib, and provide a simple and safe API in a single header file, which can be easily integrated for programming in C++ as well as other languages that can call C/C++ codes, such as R and Python.

## Methods

### The VCF and HTSlib

A VCF file consists of a header section and a body section. The header contains lines with meta-information, each starting with characters”##” or”#", while the body consists of TAB-delimited lines. A VCF file differs from a typical TAB-delimited file in several ways. Firstly, the header is too important to be ignored. A VCF that contains a tag in the body without its declaration in the header violates the specification. Secondly, the VCF can be compressed and randomly accessed using bgzip and tabix, which were originally part of SAMtools library (Li et al. 2009) but were later separated into HTSlib (Bonfield et al. 2021). Additionally, the VCF standard is established and regularly updated by the SAMtools team. Therefore, it would be more advantageous to use a specialized library, namely HTSlib, to process the VCF rather than a customized parser. HTSlib offers full functionalities to interact with the VCF, including support for BCF, format validation, compression, random access, identification of variant types, support for URL links as filenames, among others. Furthermore, HTSlib has demonstrated the best performance among many competitors (Bonfield et al. 2021). However, HTSlib was written in C language, which can be challenging for beginner programmers to use, especially when it comes to memory management.

### A C++ API

As a C++ API of HTSlib, vcfpp inherits all functionalities from HTSlib and is implemented in a single header file that can be easily integrated and used safely. There are four core classes in vcfpp, as summarized in Table 1, all of which allocate and free memory automatically and safely, enabling users to program with ease.

**Table 1.** vcfpp capabilities and implemented C++ class.

| Class | Capability |
| --- | --- |
| BcfReader | read VCF/BCF, decompression, random access, subset samples |
| BcfWriter | output VCF/BCF, compression |
| BcfRecord | access the fields, modify the fields |
| BcfHeader | access the header, modify the header |

To illustrate commonly used features and demonstrate the simplicity of vcfpp, here I showcase an example that will be used throughout the paper. In Listing 1, we count the number of heterozygous sites for each sample in a VCF file. The following code first includes a single vcfpp.h file (line 1), opens a compressed VCF file constrained to 3 samples and region”chr21” (line 2), and creates a variant record associated with the header information in the VCF file (line 3). Then it defines several types of objects to collect the results we want (line 4-5). Taking advantage of generic templates in C++, the genotype value can be of bool, char or int type so that users may control the memory consumed by their program. Then it iterates over the VCF file and processes each variant record in the loop (line 6). Here we ignore variants of other types (INDEL, SV), or if FILTER does not display”PASS", or if the QUAL value is smaller than 9 (line 8-9), though the API also allows us to do more complicated filtering should. Finally, we count the number of heterozygous variants for each diploid sample (line 10-11). The core is only 12 lines.

Listing 1. Counting the number of heterozygous genotypes for 3 samples on chr21

~~~
# include <vcfpp. h>
vcfpp : : BcfReader vcf (“your. vcf. gz”,”chr21”,”id 1, id2, id 3”) ;
vcfpp : : BcfRecord var (vcf. header) ; */ / create a variant object*
vector <int> gt ; */ / genotype can be of bool, char or int type*
vector <int> hets (vcf. nsamples, 0) ; */ / store the het counts*
while (vcf. get Next Variant (var)) {
   var. get Genotypes (gt) ;
   if (! var. isSNP () | | var .QUAL() < 9 | | var. FILTER () ! =”PASS”) continue ;
   assert (var. ploidy () == 2) ; */ / make sure it is diploid*
   for (int i = 0 ; i < vcf. nsamples ; i ++)
       hets [ i ] += abs (gt [ 2 * i + 0 ] − gt [ 2 * i + 1 ]) == 1 ;
}
~~~

## Results

To demonstrate the simplicity, portability and performance of vcfpp, the following sections include the benchmarking results and highlight the vcfppR package as an example of vcfpp working with R (R Core Team 2023).

### Working with R

While vcfpp is very simple for writing a C++ program, a single C++ header file can be easily integrated into popular script languages like R and Python. Particularly, R is designed for statistical modeling and visualization, widely used for data analyses. Therefore, I developed the vcfppR package to demonstrate how vcfpp can seamlessly work with R using Rcpp (Eddelbuettel et al. 2011). For instance, with the basic familiarity of C++ and Rcpp, we can turn the C++ code in Listing 1 into an Rcpp function to return the heterozygosity counted per sample along with the sample’s name (Listing 2), which can then be compiled and called dynamically in R using sourceCpp (Listing 3). As such, we can further process and visualize these results straightforwardly within the R ecosystem. For example, we can analyze the data by genomic region in parallel using the parallel package and then stratify the results by population, leveraging the external labels of each sample, to finally visualize them in R.

Listing 2. vcfpp-hets.cpp

~~~
# include <Rcpp. h>
# include <vcfpp. h>
using namespace Rcpp ;
*/ / [ [ Rcpp* : : *export ] ]*
List heterozygosity (std : : string vcffile,
                     std : : string region,
                     std : : string samples) {
  vcfpp : : BcfReader vcf (vcffile, region, samples) ;
  */ / here copy the l i n e s 3 −12 in l i s t i n g 1* .
  return L i s t : : create (Named(”samples”) = vcf. header. getSamples (),
                             Named(”hets”) = hets) ;
}
~~~

Listing 3. The R code compiles and calls vcfpp-hets.cpp dynamically

~~~
library (Rcpp)
sourceCpp (“vcfpp − hets. cpp”)
heterozygosity (“bcf. gz”,”chr21”,”id 1, id2, id 3”) ;
~~~

### The vcfppR package

The vcfppR package is developed and powered by the vcfpp API. To parse the VCF with vcfppR, the vcfreader and vcftable functions can rapidly read contents of the VCF into the R data types with fine control over the region, samples, variant types, FORMAT column and filters. For instance, the code in Listing 4 parses the read depth per variant (DP) in the raw called VCF by the 1000 Genomes Project through the URL link. It restricts the analysis to 3 samples with variants in”chr21:1-10000000” region of SNP type, passing the FILTER, and discarding the INFO column in the returned list. Subsequently, the visual summary can be generated by using `boxplot()` in R (see Figure 1).

**Figure 1.**
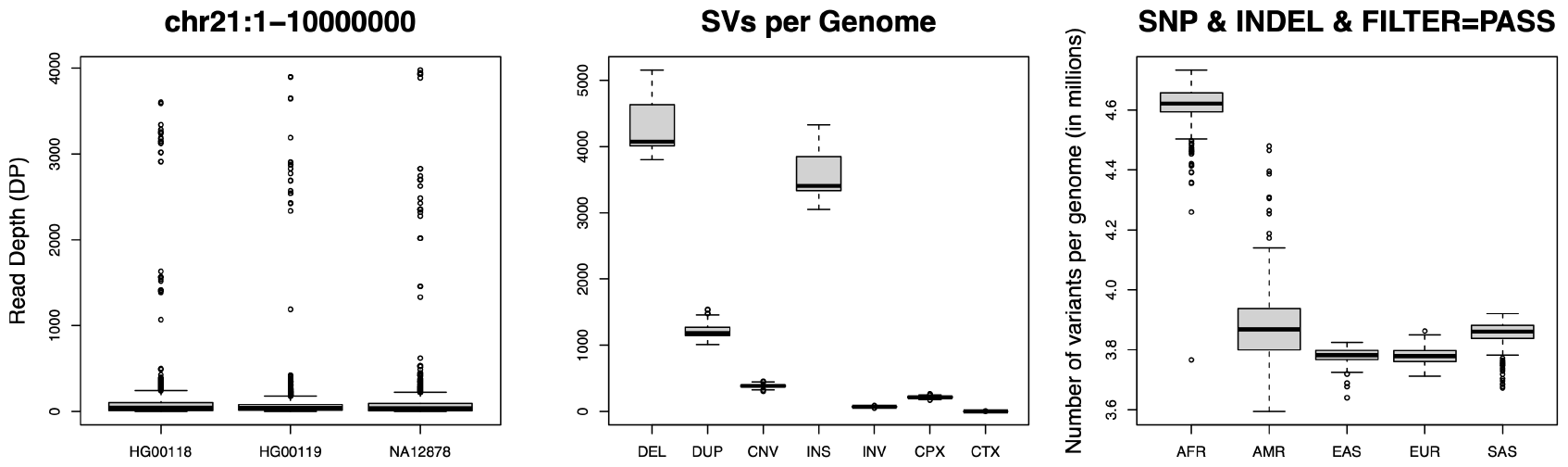
Analyzing variants discovered in the 1000 Genome Project with vcfppR (see code in supplementary).

Listing 4. Example of vcfppR::vcftable

~~~
library (vcfppR)
vcffile <-“https : // url −path”
vcf <-v c f t a b l e (v c f f i l e, region =”chr21 : 1 − 10000000”, samples=”
    NA12878, HG00118, HG00119”, format=”DP”, vartype =”snps”, pass
    = TRUE, info = FALSE)
boxplot (vcf $DP, names= vcf $samples, ylab =”Read Depth (DP)”)
~~~

Furthermore, as characterizing variants is an essential task in genomic analyses, I showcase the vcfsummary function in Figure 1, which summarizes the variants found in the latest VCF released by the 1000 Genome Project (Byrska-Bishop et al. 2022).

### Benchmarking

In addition to simplicity and portability, I showcase here the performance of vcfpp and vcfppR. For the benchmarking, I developed scripts (https://github.com/Zilong-Li/vcfpp/tree/main/scripts) to perform a common analysis of counting heterozygous genotypes per sample on a Linux server with AMD EPYC 7643 48-Core Processor. As shown in Table 2, when using the compressed (gzipped) VCF of 3202 samples and 1002753 variants, which includes only GT (genotype) in the FORMAT, the Rcpp function vcfppR::heterozygosity in Listing 2 demonstrates comparable performance to the compiled C++ code of vcfpp in Listing 1 with a minor overhead. This overhead is attributed to a list of sample names being returned to R. The dynamic script using vcfppR::vcfreader is only 1.3*×* slower than its compiled C++ counterpart, whereas the cyvcf2::VCF is 1.9*×* slower. With the streaming strategy, all scripts use little RAM, given that they only load one variant into memory at a time. However, R packages like vcfR and data.table usually load all VCF data into memory first and perform analyses later, which is referred here as the”two-step” strategy. To this end, I have also developed vcftable function in vcfppR to load the entire contents of the VCF into R for the two-step comparison. Notably, the vcfppR::vcftable is only 2.0*×* slower compared to the 19*×* slower vcfR::read.vcfR and the 119*×* slower data.table::fread. This discrepancy arises because the genotype values returned by both vcfR and data.table are characters, which are inefficient to further process in R. On the other hand, with vcfppR, an integer matrix of genotypes can be returned to R directly for fast computation. If we exclude the elapsed time of loading data, which means ignoring time marked by the *, then vcfppR demonstrates a 101× speed improvement over vcfR. Importantly, vcfpp and vcfppR offer users the full functionalities of HTSlib, including support for compressed VCF/BCF, selection of samples, regions, and variant types.

**Table 2.** Performance of counting heterozygous genotypes per sample with VCF of 3202 samples and 1002753 variants. * used by loading data in two-step strategy.

| API/Function | Time (s) | Ratio | RAM (Gb) | Strategy |
| --- | --- | --- | --- | --- |
| vcfpp | 97 | 1.0 | 0.006 | streaming |
| vcfppR::heterozygosity | 109 | 1.1 | 0.074 | streaming |
| vcfppR::vcfreader | 150 | 1.3 | 0.136 | streaming |
| cyvcf2::VCF | 212 | 1.9 | 0.036 | streaming |
| vcfppR::vcftable | 206*+12 | 2.0 | 64.7 | two-step |
| vcfR::read.vcfR | 625*+1219 | 19.0 | 97.5 | two-step |
| data.table::fread | 313*+11243 | 119.1 | 77.3 | two-step |

## Discussion

Here I have developed vcfpp, a fast and flexible C++ API for high-performance genetic variant analyses with the VCF/BCF input. Its simplicity and portability can be very valuable for both developing packages and writing in-house scripts. The vcfppR package is a great example of vcfpp working with R, and there are also examples of developing a Python API for vcfpp available on GitHub. In vcfppR, there are vcfreader and vcftable functions that can process the variants. The vcftable is specifically designed for reading the VCF into the tabular data structure in R, but it can only read a single FORMAT item in one pass over the VCF. In contrast, vcfreader serves as the full R-bindings of vcfpp and allows iterative parsing of variants, giving users the flexibility to decide the information to retrieve for each variant. As such, many packages written in C++ using a customized VCF parser can be simply replaced with vcfpp to offer more functionalities. For instance, vcfpp can be found successfully implemented in the imputation software programs STITCH (Davies, Flint, et al. 2016) and QUILT (Davies, Kucka, et al. 2021) to parse large reference panels in the compressed VCF/BCF.

### Software and Code

The latest release of vcfpp.h and documentation can be found at https://github.com/Zilong-Li/vcfpp. The vcfppR package can be installed through CRAN for all platforms (https://CRAN.R-project.org/package=vcfppR). Scripts for the benchmarking are available at https://github.com/Zilong-Li/vcfpp/tree/main/scripts.

## Acknowledgments

I would like to thank Anders Albrechtsen at Copenhagen University and Robert W Davies at Oxford University for their helpful comments. They are statisticians as well as R enthusiasts working on genetics, whom I work with and have learned a lot from. Also, I want to thank Cindy G. Santander and the reviewers for helping me to improve the quality of this article.

## Supplementary

Listing 5. R code produces Figure 1. Vcffiles (not shown) are url links.

~~~
library (vcfppR)
par (mfrow=c (1, 3)) *## l ayout in p l o t s*
vcf <-v c f t a b l e (v c f f i l e, region =”chr21 : 1 − 10000000”, samples=”
    NA12878, HG00118, HG00119”, format=”DP”, vartype =”snps”, pass
    = TRUE, info = FALSE)
boxplot (vcf $DP, names= vcf $samples, ylab =”Read Depth (DP)”)
svfile <-“https : // ftp. 1000 genomes. ebi. ac. uk/ vol 1 / ftp / data _
    collections / 1000G_ 2504 _high_coverage/ working/ 20210124. SV_
    Illumina _ In t eg ra t i o n / 1KGP_ 3202. gatksv _ s v t o o l s _ novelins .
    freeze _V3 .wAF. vcf. gz”
sv <-vcfsummary (svfile, svtype = TRUE)
boxplot (sv [ c (“DEL”,”DUP”,”CNV”,”INS”,”INV”,”CPX”,”CTX”) ],
         main =”SVs per Genome s t r a t i f i e d by SV types”)
ped <-read. table (“https : // f t p. 1 0 0 0 genomes. ebi. ac. uk/ vol 1 / f t p /
     data _ c o l l e c t i o n s / 1000G_ 2504 _high_coverage/ 20130606 _g1k_ 3202 _
     samples_ped_population. t x t”, h=T)
ped <-ped [ order (ped$ Superpopulation), ]
supers <-unique (ped$ Superpopulation)
all <-p a r a l l e l : : mclapply (v c f f i l e s, vcfsummary, pass = TRUE, mc.     cores = 23)
samples <-all [ [ 1 ] ] $samples
snps <-Reduce (“+”, lapply (all,”[ [”,”SNP”))
ind els <-Reduce (“+”, lapply (all,”[ [”,”INDEL”))
o <-sapply (supers, function (pop) {
   id <-subset (ped, Superpopulation == pop) [,”SampleID”]
   ord <-match (id, samples)
   ( snps [ ord ] + ind els [ ord ]) / 1 e6
})
boxplot (o, main =”SNP & INDEL with FILTER=PASS”, ylab =”
   Number of variants per genome (in millions)”)
~~~

